# Breathing Behaviors in Common Marmoset (*Callithrix jacchus*)

**DOI:** 10.1101/2020.07.27.223990

**Authors:** Mitchell Bishop, Ariana Turk, Afuh Adeck, Shahriar SheikhBahaei

## Abstract

The respiratory system maintains homeostatic levels of oxygen (O_2_) and carbon dioxide (CO_2_) in the body through rapid and efficient regulation of frequency and depth (tidal volume) of breathing. Many studies on homeostatic control of breathing have been done on rodent animal models, in which they have relatively reduced chemosensitivities when compared with primates. The use of common marmoset (*Callithrix jacchus*), a New World non-human primate model, in neuroscience is increasing, however, the data on their breathing is limited and their respiratory behaviors have yet to be characterized. Using Whole-body Plethysmography in room air as well as in hypoxic (low O_2_) and hypercapnic (high CO_2_) conditions, we defined breathing behaviors in awake, freely behaving marmosets. Additionally, we optimized an analysis toolkit for unsupervised analysis of respiratory activities in common marmoset. Our data indicate that marmosets’ exposure to hypoxia decreased metabolic rate and increased sigh rate. However, the hypoxic condition did not augment the ventilatory response. Hypercapnia, on the other hand, increased both the frequency and tidal volume. In this study, we described breathing behaviors of common marmosets in a variety of O_2_ and CO_2_ conditions.

## Introduction

Mammals rely on fresh and continuous supply of oxygen (O_2_) from the environment and efficient removal of carbon dioxide (CO_2_) and other metabolic waste products from their body. The intricate respiratory system ensures the homeostatic state of the arterial partial pressure of O_2_ (*P*O_2_) and CO_2_ (*P*CO_2_) in the blood by executing rhythmic movement of the respiratory pump, which include the intercostals and the diaphragm muscles (Del Negro et al. 2018). The inception of this respiratory rhythm occurs within the preBötzinger complex (preBötC), a functionally specialized region in the ventrolateral medulla of the brainstem (Smith et al. 1991; Del Negro et al. 2018). Activities of the preBötC are modulated by specialized peripheral and central chemosensors that adjust the respiratory drive to regulate homeostatic levels of *P*O_2_ and *P*CO_2_ (Heymans and Bouckaert 1930; O’Regan and Majcherczyk 1982; Guyenet 2014; Sheikhbahaei et al. 2018; Angelova et al. 2015; van der Heijden and Zoghbi 2019; Guyenet et al. 2019; Sheikhbahaei et al. 2017; Del Negro et al. 2018).

Studies on homeostatic control of breathing have been done mainly on rodent models, in which the experiments are mostly performed during the day i.e., rodent’s normal inactive period. Since, in general, rodents have relatively reduced chemosensitivities compared with primates (Hazari and Farraj 2015), there is little assurance on if these results can effectively be extrapolated to humans. Therefore, use of non-human primates (NHPs) has been proposed to fill this gap and translate rodent data to humans (Sheikhbahaei 2020). The common marmoset (*Callithrix jacchus*) is a New World NHP with a small body size (250 – 600 g) similar to a rat. Ease of handling, high reproductive efficacy, and lack of zoonotic risks compared to Old World NHPs make marmosets an attractive and powerful NHP model for biomedical and neuroscience research (Abbott et al. 2003). Recently, the use of marmosets in neuroscience research settings has increased as the marmoset has been proposed as a primate model to study behavioral neuroscience, including vocalization (Prins et al. 2017; Miller et al. 2016; Walker et al. 2017). Although breathing is essential for vocal production, the basic characteristics of breathing behaviors in marmoset are not yet defined.

Here, we used Whole-body Plethysmography to record breathing behaviors of awake, freely behaving common marmosets in room air, as well as during hypoxic (low inspired O_2_) and hypercapnic (high inspired CO_2_) conditions. We also modified and optimized an analysis toolkit for automated characterization of respiratory indices in laboratory animals. We found that while exposure of marmosets to hypoxia increased sigh rate and decreased overall animal metabolic rate, the hypoxia-induced augmentation of ventilatory response was diminished. On the other hand, hypercapnic conditions increased both frequency and depth of breathing similar to other mammals of similar size.

## Material and Methods

### Animals

We used sixteen common marmosets (*Callithrix jacchus*) (8 males, 8 females; 394 ± 5 g; 40 ± 1 months) for measuring and defining breathing behaviors. All experiments were performed in accordance with the National Institutes of Health Guide for the Care and Use of Laboratory Animals and were approved by the Animal Care and Use Committee of the Intramural Research Program of the National Institute of Mental Health. Animals were housed in temperature-controlled facilities on a normal light-dark cycle (12h:12h, lights on at 7:00 AM). They lived in paired or family-grouped housing and were given tap water *ad libitum*.

### Measurement of marmoset respiratory activity

Marmoset respiratory activity was measured using Whole-body Plethysmography. Awake animals were placed in the Plexiglas chamber (∼3 L) which was flushed with 21% O_2_, 79% N_2_, 22-24 °C, at a rate of 1.2 L /min during measurements of baseline respiratory behavior (Figure 1). Concentrations of O_2_ and CO_2_ in the chamber were monitored using a fast-response O_2_ / CO_2_ analyzer (ML206, AD Instruments). All experiments were performed at the same time of day (between 10:00 and 14:00 hours) to account for possible circadian changes in base level physiology (Iizuka et al. 2010). For measuring the respiratory behaviors during hypoxia, following a 40-minute baseline period, the chamber was flushed with 10% O_2_, 90% N_2_, 22-24 °C, at a rate of 1.2 l min^-1^. After 10 minutes of exposure to hypoxic conditions, the gas concentration in the chamber was changed to room air for another 10 minutes (Figure 1 – figure supplement 1). Marmoset respiratory activity was also measured during exposure to hypercapnic conditions. Following a 40-minute baseline period, the chamber was flushed with 6% CO_2_, 60% O_2_, 34% N_2_, 22-24 °C, at a rate of 1.2 l min^-1^. After 10 minutes of exposure to hypercapnic conditions, the chamber was then flushed with room air for another 10 minutes. Hyperoxic condition (60% O_2_) was used to prevent any hypoxia associated with hypercapnia as used routinely in rodents (Teppema et al. 1997; Sheikhbahaei et al. 2018). Respiratory data were acquired with Power1401 (CED; RRID: SCR_017282) interface and transferred to Spike2 software (CED; RRID: SCR_000903). To prevent any acclimatization confound, each animal was only placed once in the plethysmography chamber and randomly assigned to either hypoxia or hypercapnia experiment.

**Figure 1.**
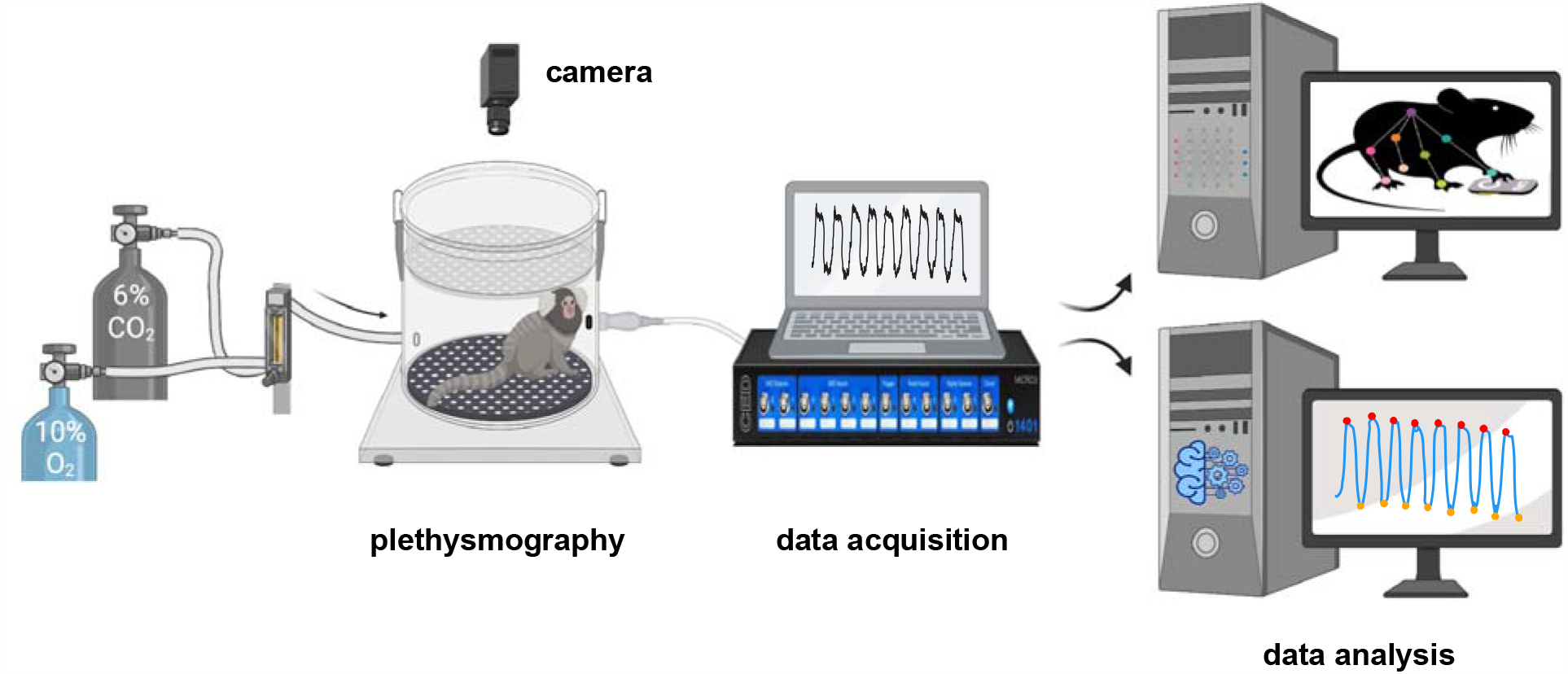
Experimental pipeline for measurement and analysis of marmoset respiratory behaviors. After a 40-minute baseline period at room air (21% O_2_, ∼0% CO_2_, and 79% N_2_), the breathing behavior of animal was studied under either hypoxic (10% O_2_; 10 min) or hypercapnic (6% CO_2_; 10 min) conditions. Raw respiratory signal is later cleaned and analyzed offline (see Methods for details). Video of spontaneous activity in the chamber at baseline and during each challenge were used to train a DeepLabCut model to track the animal’s body.

### Calculation of metabolic rate

For measuring metabolic rate (M_R_), we calculated CO_2_ production using the following equation and expressed as percent: M_R_ = •CO_2_ x F_R_ / body mass, where •CO _2_ is the peak changes in the [CO_2_] in the chamber as measured by the gas analyzer. F_R_ is the flow rate through the plethysmography chamber (i.e., 1.2 l min^-1^), and body mass is marmoset body mass (g).

### Automated quantification of animal activity

We tracked 10 points on the marmoset head and body (*n* = 3 animal per challenge) from an overhead view of the plethysmograph using WhiteMatter e3Vision cameras (e3Vision camera; e3Vision hub; White Matter LLC). We used DeepLabCut version 2.10.2 for pose estimation of these features (Nath et al. 2019; Mathis et al. 2018). We labeled 656 total frames from 16, 20–30-minute videos recorded at 60 fps (95% was used for model training). We used ResNet-50-based neural network with default parameters for 4 iterations with 5 shuffles, and the test error was: 29.8 pixels, train: 2.4 pixels, with 0.6 p-cutoff, test error was: 14.0 pixels, train: 2.4 pixels (image size 600 by 800 pixels).

Missing feature coordinates were then filled using methods from the B-SOiD Python toolkit (Hsu and Yttri 2019). We used the average position of 5 points on the head for further analysis after qualitative assessment of consistent labeling accuracy. By dividing the labeled images in quadrants along the X- and Y-axes (X = 400 pixels, Y=300 pixels), we counted the number of times large changes in position (i.e., movement) occurred. Quadrant positions were down sampled to 2 seconds to avoid counting quadrant changes from when the animal paused near the dividing lines. Additionally, successive Euclidean distances were calculated for each point across each frame of the videos to produce total movement. Total movement was then divided by length of condition in minutes to obtain rate of activity in each condition.

### Respiratory data analysis

All animals in the study were included in the analysis. Plethysmography data were imported to Python using Neo Python package (Van Rossum and Drake 2011; Garcia et al. 2014). We wrote a custom Python script using methods from Neurokit2, NumPy, and Pandas software packages (McKinney 2010; van der Walt et al. 2011; Makowski et al. 2020). Areas of the signal with frequencies above 300 cycles per minute (∼3.3 Hz) were excluded from analysis, as they were likely artifact resulting from movement inside the chamber. To ensure that we capture the full change in ventilation, we used steady-state responses to hypoxia and hypercapnia and analyzed the data 5 minutes after the start of each challenge. Neurokit2 methods were used for signal cleaning and extraction of instantaneous frequency, T_TOT_ (total time of breath), T_I_ (time of inspiration), T_E_ (time of expiration), and amplitude [i.e., tidal volume (*V*_*T*_)] from trough to peak of the signals (see Figure 2). The calculated *V*_T_ was normalized to the body mass (g) of each animal. Mean inspiratory flow rate was defined as the ratio of *V*_T_ to T_I_ and used as an index of inspiratory drive (R_D_). During hypoxia and hypercapnia challenges, the respiratory signals were analyzed in 1-minute epochs to consider local changes in respiration parameters.

**Figure 2.**
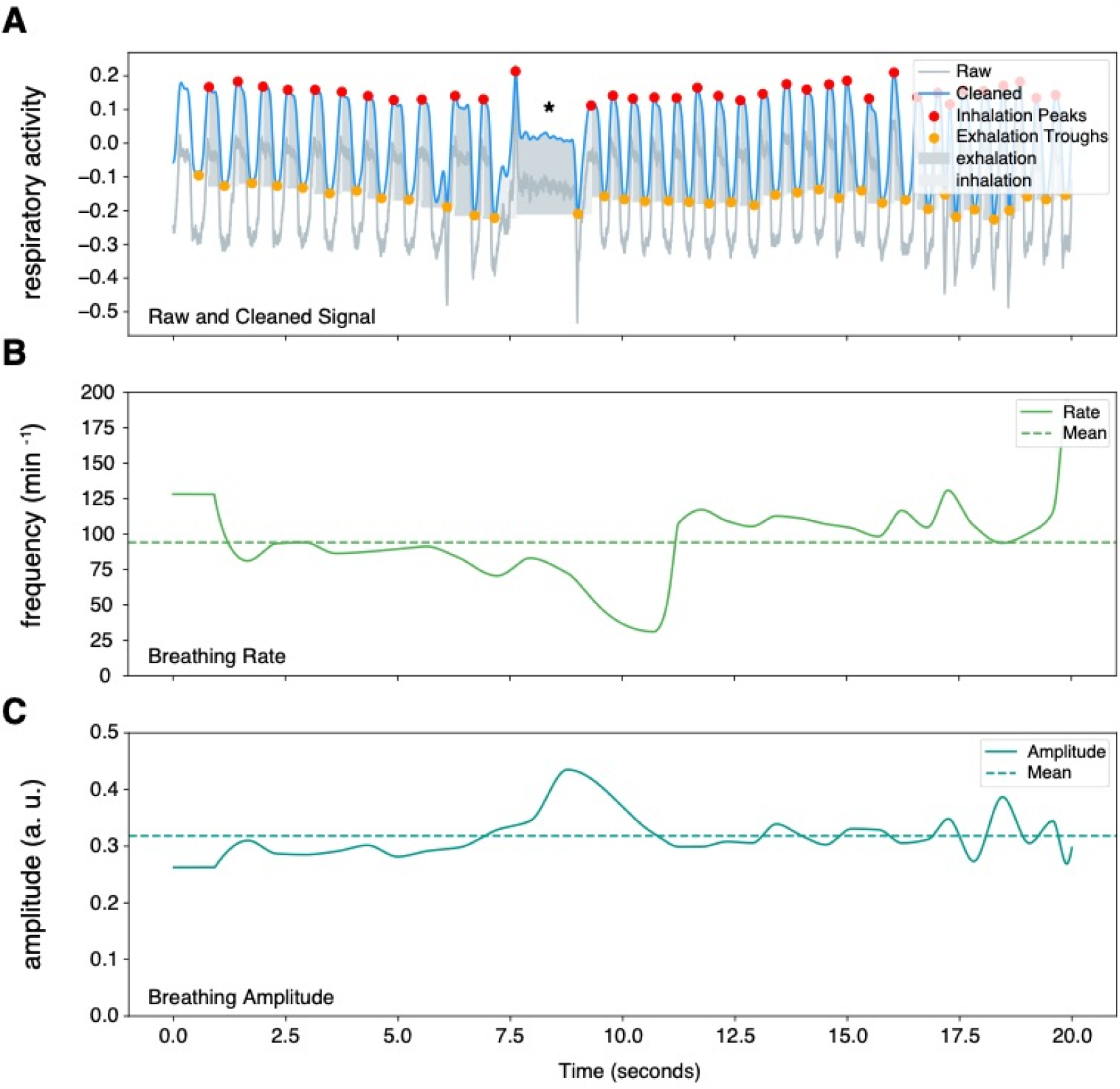
Sample plot output from NeuroKit2 Python package. Representative respiratory trace is sampled from a single male marmoset during hypercapnia challenge. (**A**) NeuroKit2 was used for signal detrending and smoothing, peak and trough extraction, as well as respiratory phase. (**B & C**) We also used NeuroKit2 methods for instantaneous measurement of breathing frequency (*f*_R_) (**B**) and breathing amplitude (*V*_T_) (**C**). This sample also contained respiratory changes during a *phee* call (marked by *). a. u. – arbitrary unit.

High frequency breathing (i.e., sniffing) was defined as any breathing frequencies between 250 cycles (2.5 Hz) and 300 cycles per minute. Apneas were defined by breathing cycles with T_TOT_ greater than 3 times the average for each animal. Augmented breaths (i.e., sighs) were readily identifiable by using the criteria described in rats (Sheikhbahaei et al. 2018; Sheikhbahaei et al. 2017) and measured during the baseline and experimental conditions.

Two measures of rate variability were also calculated as described elsewhere (Soni and Muniyandi 2019). SD1 is a measure of dispersion of T_TOT_ perpendicular to the line of identity in the Poincaré plots, therefore demonstrating short term variability. SD2 is a measure of dispersion of T_TOT_ along the line of identity in the Poincaré plots, demonstrating long term variability in respiratory rate.

SD1 and SD2 are calculated by:

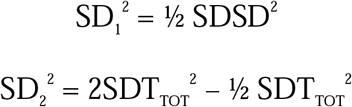

Where SDSD is the standard deviation of successive differences in T_TOT_ and SDT_TOT_ is the standard deviation in T_TOT_.

All data were tested with Shapiro-Wilk test for normality and statistically compared by Wilcoxon matched-pairs signed rank test or Mann–Whitney *U* rank test as appropriate in Prism 9 (Graphpad, Inc; RRID: SCR_002798). Data is reported as mean ± SEM.

### Data Availability

All the code is available on the NGSC GitHub (https://github.com/NGSC-NINDS/Marm_Breathing_Bishop_et_al_2021). The data generated in Figures 3 – 10 are provided in the source files.

**Figure 3.**
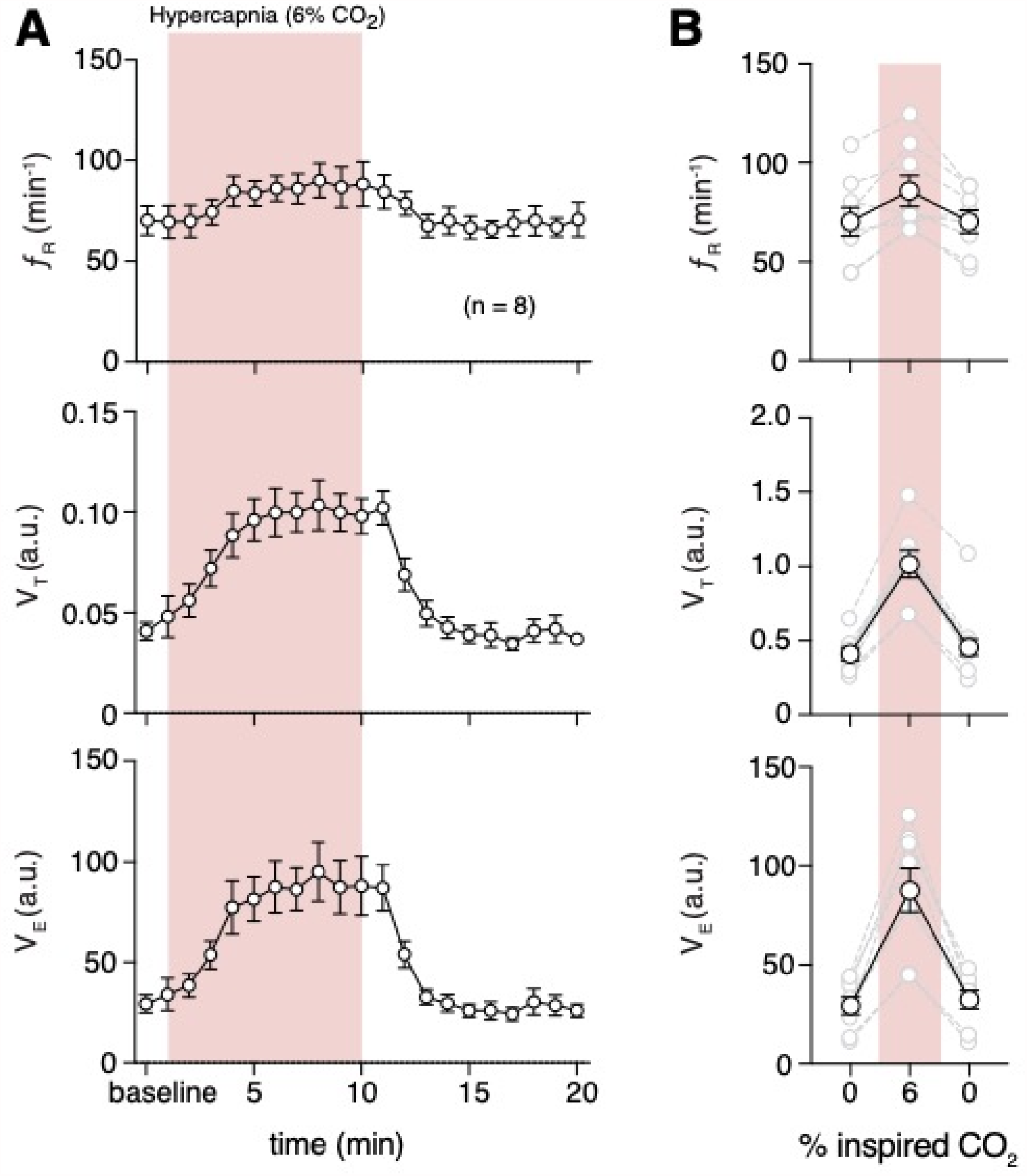
Hypercapnia challenge induced changes in respiratory behavior. (**A**) Measurements of breathing rate (*f*_R_), tidal volume (*V*_T_), and minute ventilation (*V*_E_) were averaged across 1-minute epochs for assessment of local changes in each parameter. (**B**) Summaries of each feature at baseline, following 5 minutes exposure to hypercapnia, and in the 5 minutes immediately following the end of challenge. We observed increases in respiratory frequency (*p* = 0.023, Wilcoxon matched-pairs signed rank test), *V*_*T*_ (*p* = 0.008, Wilcoxon matched-pairs signed rank test), and *V*_*E*_ (*p* = 0.008, Wilcoxon matched-pairs signed rank test) during hypercapnia. Data are shown as individual (gray lines) and mean (black line) values ± SEM. a. u. – arbitrary unit.

## Results

### Validation of the Neurokit2 as an analysis toolkit in experimental animal models

To analyze resting rate of breathing (*f*_R_), tidal volume (*V*_T_), and minute ventilation (*V*_E_), we first benchmarked the Neurokit2 analysis toolkit against a conventional method of analyzing respiratory data in conscious mammals (Sheikhbahaei et al. 2018; Sheikhbahaei et al. 2017). We did not identify any differences in values of *f*_R_, *V*_T_, and *V*_E_ when using Neurokit2 or conventional methods (*n* = 3) (Figure 2 – figure supplement 1).

### Resting respiratory behavior in adult marmosets

The *f*_R_ at room air (normoxia /normocapnia) calculated from breath-to-breath time (T _TOT_) was similar in female (79 ± 7 breaths min^-1^, *n* = 8) and male (78 ± 8 breaths min^-1^, *n* = 8) adult marmosets (*p* = 0.88, Mann-Whitney test) (Figure 2 – figure supplement 2). The *V*_T_, calculated from trough to peak amplitude and normalized to body mass, was similar in female (0.43 ± .10 a.u.) and male (0.53 ± 0.08 a.u.) adult marmosets as well (*p* = 0.37, Mann-Whitney test). Additionally, baseline *V*_E_ was similar in female (35 ± 10 a.u.) and male (42 ± 8 a.u.) marmosets (*p* = 0.38, Mann-Whitney test). Two marmosets (1 male and 1 female) showed prolonged breath holding (11 ± 2 breaths hr^-1^ for 4.3 ± .1 sec).

### Hypercapnic Ventilatory Response

We also measured changes in *f*_R_, *V*_T_, and *V*_E_ before, during, and after hypercapnic challenge (6% CO_2_ in the inspired air). The magnitude of change in *f*_R_, *V*_T_, and *V*_E_, were similar between females and males during hypercapnia (*n* = 4 per sex, Figure 3 – figure supplement 1), so we grouped them for further analyses. Increasing CO_2_ inside the chamber increased *f*_R_ (87 ± 8 vs 74 ± 8 breaths min^-1^ in baseline, *p* = 0.039, Wilcoxon matched-pairs signed rank test), *V*_T_ (1.04 ± .11 vs 0.4 ± 0.05 a.u. in baseline, *p* = 0.008, Wilcoxon matched-pairs signed rank test) and *V*_E_ (81 ± 11 vs 32 ± 5 a.u. in baseline, *p =* 0.008, Wilcoxon matched-pairs signed rank test) (Figure 3).

Hypercapnic-induced increase in *f*_R_ was mainly due to decrease in T_I_ (0.26 ± 0.02 vs 0.36 ± 0.03 sec at baseline, *p* = 0.008, Wilcoxon matched-pairs signed rank test) rather than T_E_ (0.48 ± 0.06 vs 0.58 ± 0.09 at baseline sec, *p* = 0.078, Wilcoxon matched-pairs signed rank test). As expected, R_D_ was also increased (4.1 ± 0.4 vs 1.2 ± 0.2 a.u. in baseline, *p* = 0.008, Wilcoxon matched-pairs signed rank test) during hypercapnia (Figure 4 & Figure 4 – figure supplement 1).

**Figure 4.**
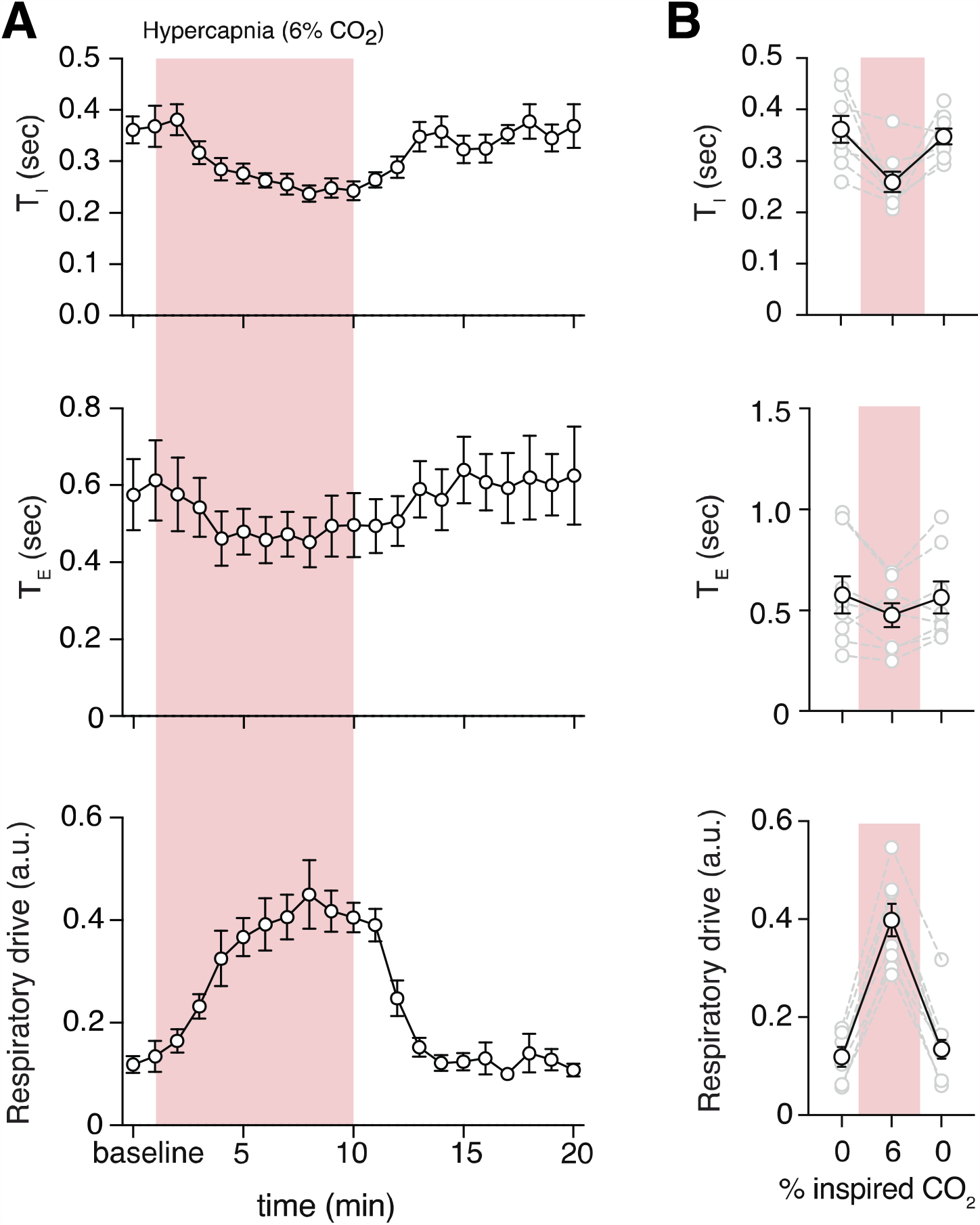
Hypercapnia induced changes in respiratory features. (**A**) Measurements of inspiratory time (T_I_), expiratory time (T_E_), and respiratory drive (R_D_) were averaged across 1-minute epochs for assessment of local changes in each parameter. (**B**) Summaries of each feature at baseline (0% inspired CO_2_), following 5 minutes exposure to 6% hypercapnia, and in the first 5-minutes following the end of hypercapnic challenge. During hypercapnia, we observed decreases in T_I_ (*p* = 0.008, Wilcoxon matched-pairs signed rank test), T_E_ (*p* = 0.078, Wilcoxon matched-pairs signed rank test), and increase in R_D_ (*p* = 0.008, Wilcoxon matched-pairs signed rank test) Data are shown as individual (gray lines) and mean (black line) values ± SEM. a. u. – arbitrary unit.

Subsequently, we measured regularity of respiration via cycle-to-cycle dispersion of T_TOT_ in baseline and hypercapnic condition as shown in Poincaré plots (Figure 5). We quantified the regularity of breathing (Sheikhbahaei et al. 2017) by SD1 and SD2 [see Methods and (Soni and Muniyandi 2019)]. The baseline SD1 and SD2 were greater than those during hypercapnia (132 ± 17 vs 550 ± 115 a.u. in baseline, *p* = 0.008, and 198 ± 34 vs 758 ± 138 in baseline, *p* = 0.008, respectively; Wilcoxon matched-pairs signed rank test) (Figure 5 and Figure 5 – figure supplement 1).

**Figure 5.**
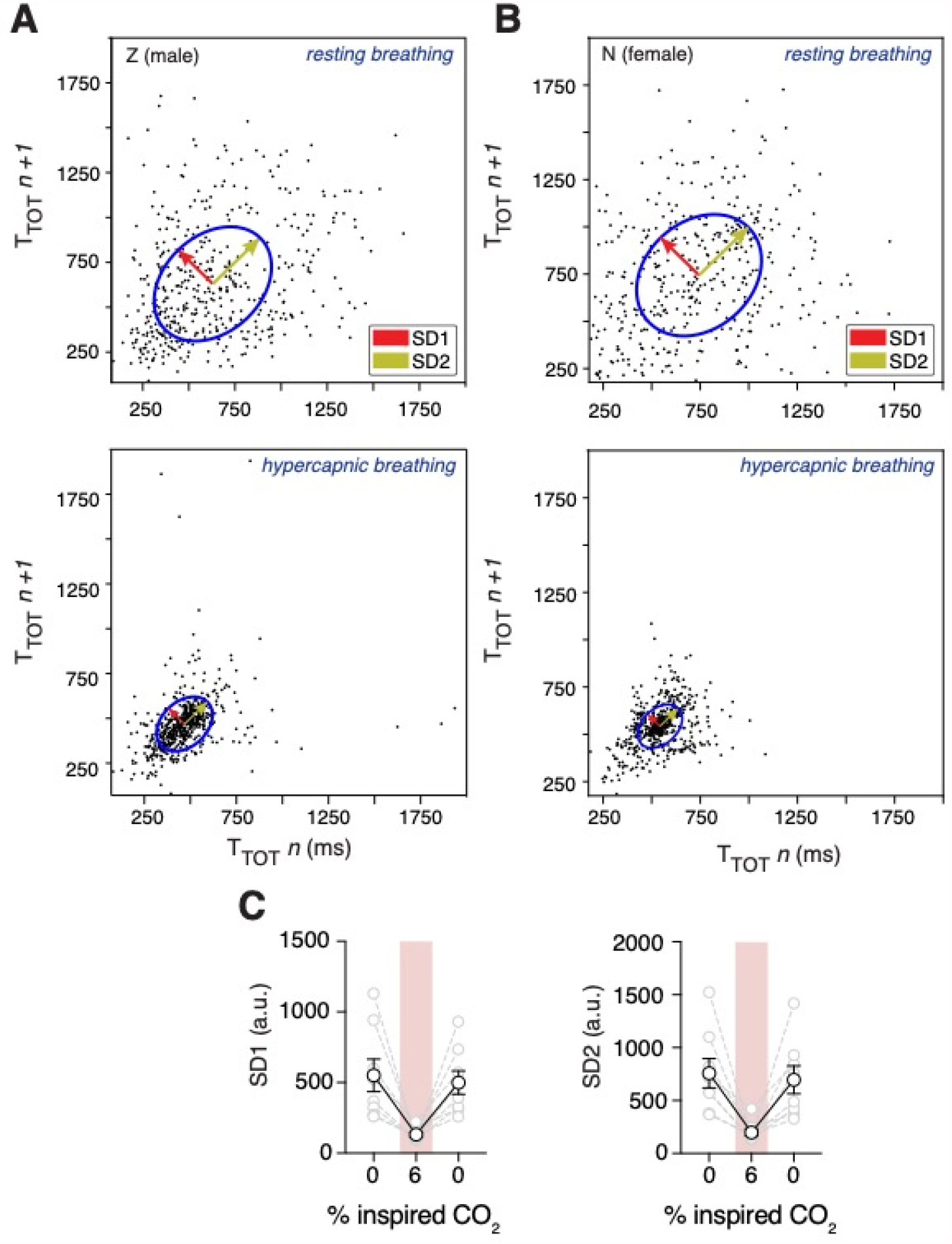
Changes in respiratory rate variability following hypercapnia challenge. Representative Poincaré plots of total cycle duration (T_TOT_) for the n^th^ cycle versus T_TOT_ for the n^th^+1 cycle during baseline (room air) and hypercapnic (6% CO_2_) conditions in male (**A**) and female (**B**) marmosets. (**C**) Grouped data illustrating changes in SD1 and SD2 before, during, and after hypercapnia challenge. Respiratory rate variability decreased in both measures during hypercapnia compared to baseline (SD1: *p* = 0.008; SD2: *p* = 0.008; Wilcoxon matched-pairs signed rank test). Data are shown as individual (gray lines) and mean (black line) values ± SEM. a. u. – arbitrary unit.

### Hypoxic Ventilatory Response

We then measured changes of *f*_R_, *V*_T_, and *V*_E_ during systemic hypoxic challenges (10% O_2_ in the inspired air) with respect to the baseline. Similar to hypoxia, the magnitude of the change in *f*_R_, *V*_T_, and *V*_E_ were not different in females and males during hypoxia (*n* = 4 per sex) (Figure 6 – figure supplement 1), therefore we combined all the data from both sexes. In the first minute of the hypoxic challenge, *V*_T_ and *V*_E_ increased by 17 ± 12% and 17 ± 14%, respectively (Figure 6). This initial increase in ventilation may be due to hypoxic induced carotid bodies activation. However, changing the inspired O_2_ from 21% (room air) to 10% did not elicit overall changes in *f*_R_ (74 ± 5 vs 82 ± 7 breaths min^-1^ in baseline, *p* = 0.3, Wilcoxon matched-pairs signed rank test) or *V*_E_ (29 ± 6 vs 46 ± 12 a.u. in baseline, *p* = 0.11, Wilcoxon matched-pairs signed rank test), but decreased *V*_T_ (0.39 ± 0.08 vs 0.54 ± .11 a.u. in baseline, *p* = 0.078, Wilcoxon matched-pairs signed rank test) (Figure 6).

**Figure 6.**
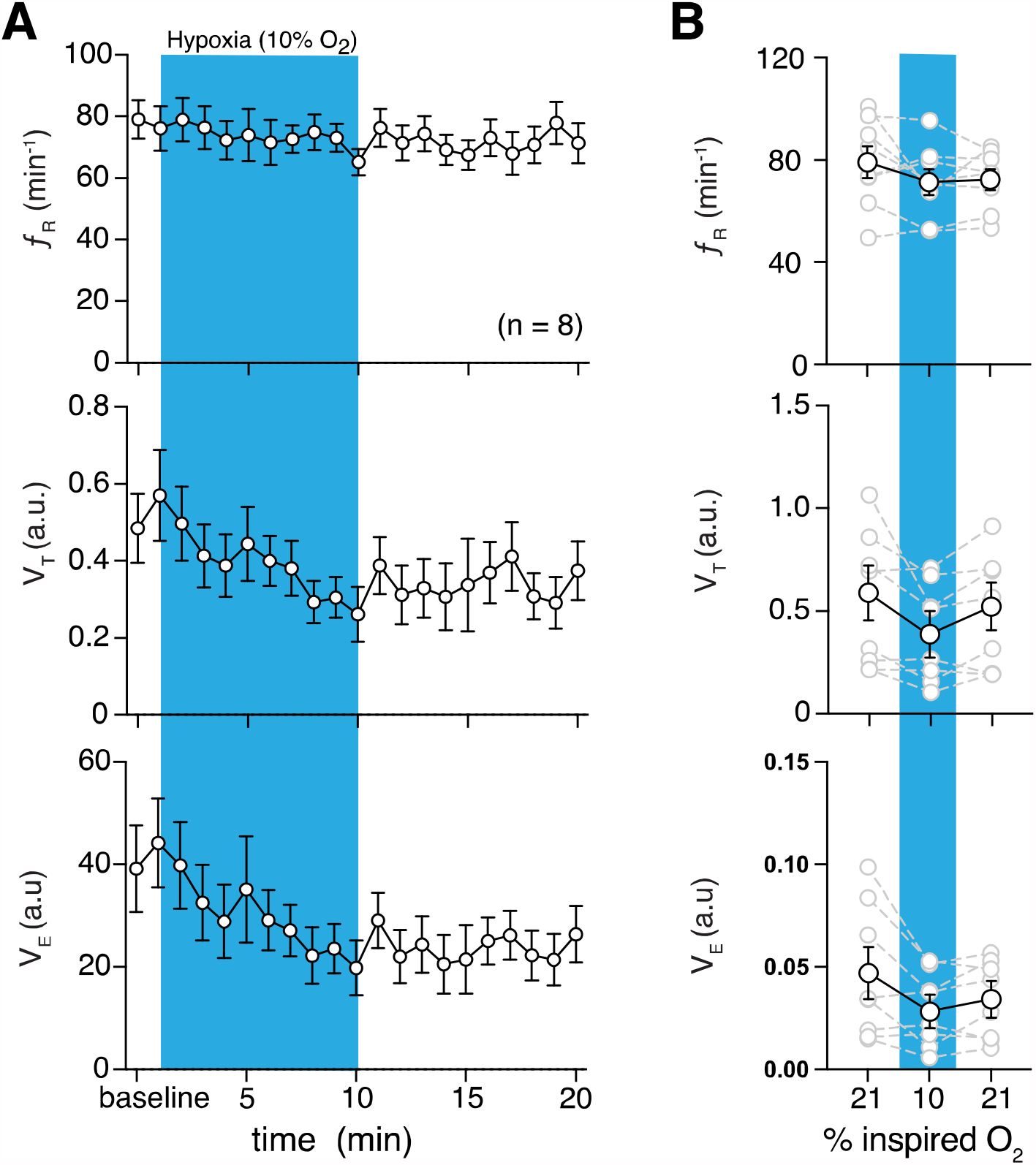
Hypoxic challenge induced changes in respiratory behavior. (**A**) Measurements of breathing rate (*f*_R_), tidal volume (*V*_T_), and minute ventilation (*V*_E_) were averaged across 1-minute epochs for assessment of local changes in each parameter. (**B**) Summaries of each feature at baseline, following 5 minutes exposure to hypoxic (10% O_2_) challenge, and in the 5 minutes immediately following the end of challenge. During hypoxia challenge we saw no changes in respiratory frequency (*p* = 0.31, Wilcoxon matched-pairs signed rank test) and *V*_E_ (*p* = 0.11, Wilcoxon matched-pairs signed rank test) compared to baseline. *V*_T_ decreased during hypoxia challenge (*p* = 0.078, Wilcoxon matched-pairs signed rank test). Immediately following the challenge, we saw no changes in respiratory frequency (*p* = 0.11, Wilcoxon matched-pairs signed rank test) compared to baseline, and a post-challenge decrease in V_T_ (*p* = 0.078, Wilcoxon matched-pairs signed rank test) and V_E_ (*p* = 0.078, Wilcoxon matched-pairs signed rank test) compared to baseline. Data are shown as individual (gray lines) and mean (black line) values ± SEM. a. u. – arbitrary unit.

Then, we also calculate changes in time of inspiration (T_I_), time of expiration (T_E_), and respiratory drive (R_D_) during hypoxic challenge with respect to baseline. Since T_I_, T_E_, and R_D_ were not different in females and males during hypoxia (*n* = 4 per sex) (Figure 7 – figure supplement 1), we combined their data. While hypoxia did not change T_I_ (0.34 ± 0.02 vs 0.30 ± 0.02 sec in baseline, *p* = 0.46, Wilcoxon matched-pairs signed rank test) and T_E_ (0.55 ± 0.1 vs 0.50 ± 0.1 sec in baseline, *p* = 0.4, Wilcoxon matched-pairs signed rank test), R_D_ was decreased during hypoxia (12 ± 2 vs 16 ± 3 a.u. in baseline, *p* = 0.008, Wilcoxon matched-pairs signed rank test) after 5 minutes of challenge (Figure 7).

**Figure 7.**
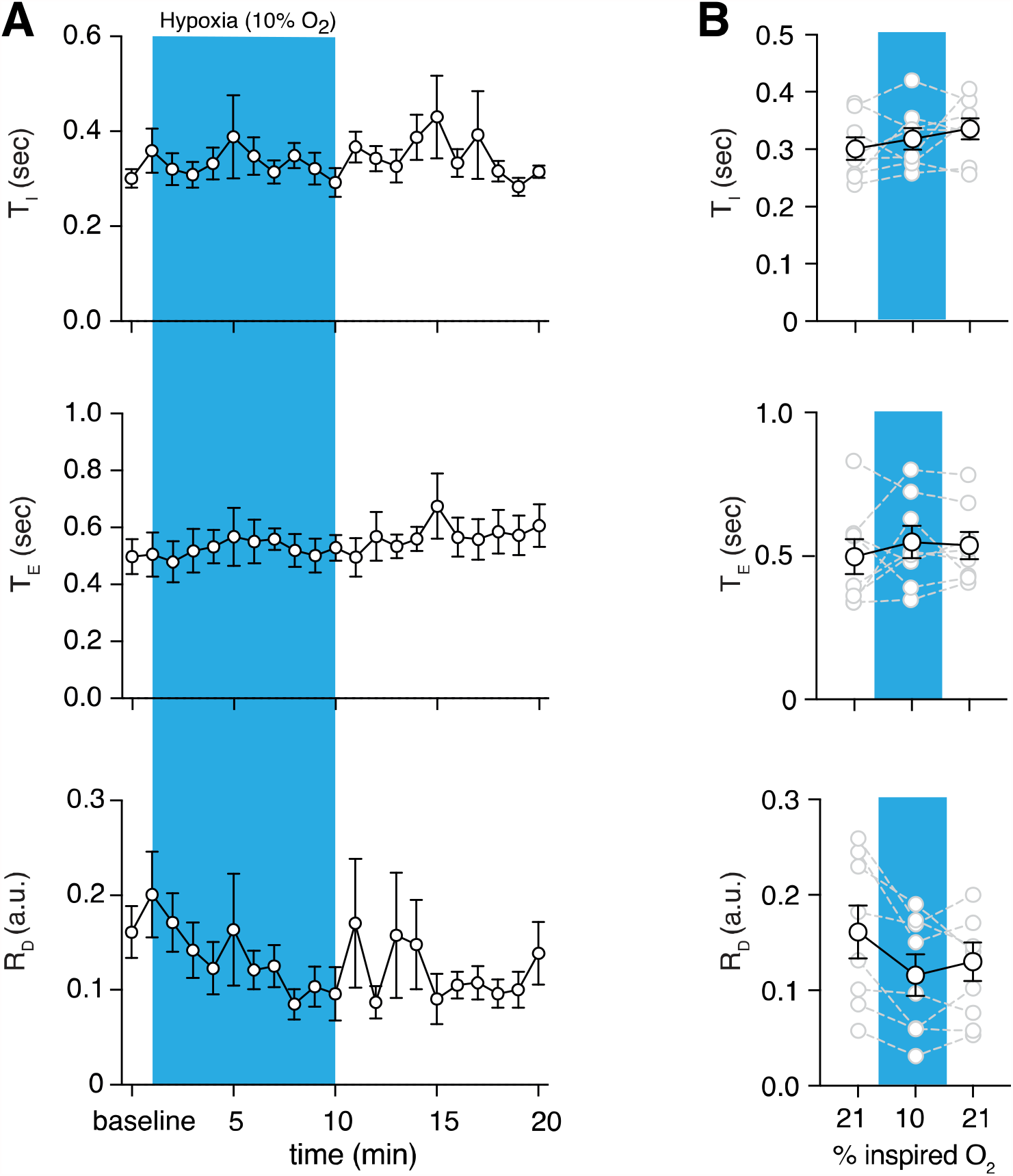
Hypoxic and post-hypoxic challenge induced changes in respiratory features. (**A**) Measurements of inspiratory time (T_I_), expiratory time (T_E_), and respiratory drive (R_D_) were averaged across 1-minute epochs for assessment of local changes in each parameter. (**B**) Summaries of each feature at baseline, following 5 minutes exposure to challenge until end of challenge, and in the 5 minutes immediately following the end of challenge. During hypoxic challenge, we saw no changes in respiratory T_I_ (*p* = 0.5) or T_E_ (*p* = 0.4), but decrease in R_D_ during (*p* = 0.008, Wilcoxon matched-pairs signed rank test) compared to baseline. We did observe post-hypoxic challenge increase in T_I_ (*p* = 0.055) and R_D_ (*p* = 0.023) and no change in T_E_ (*p* = 0.46, Wilcoxon matched-pairs signed rank test). Data are shown as individual (gray lines) and mean (black line) values ± SEM. a. u. – arbitrary unit.

We also measured the effects of hypoxia on regularity of breathing (Figure 8). We combined the data from male and female marmosets as there were no sex differences when we measured irregularity of breathing (Figure 8 – figure supplement 1). We quantified the regularity of breathing by generating Poincaré plots and measuring SD1 and SD2. The baseline SD1 and SD2 were similar during hypoxia (330 ± 39 vs 374 ± 42 in baseline, *p* = 0.4, and 419 ± 52 vs 488 ± 68 in baseline, *p* = 0.6, respectively; Wilcoxon matched-pairs signed rank test) (Figure 8).

**Figure 8.**
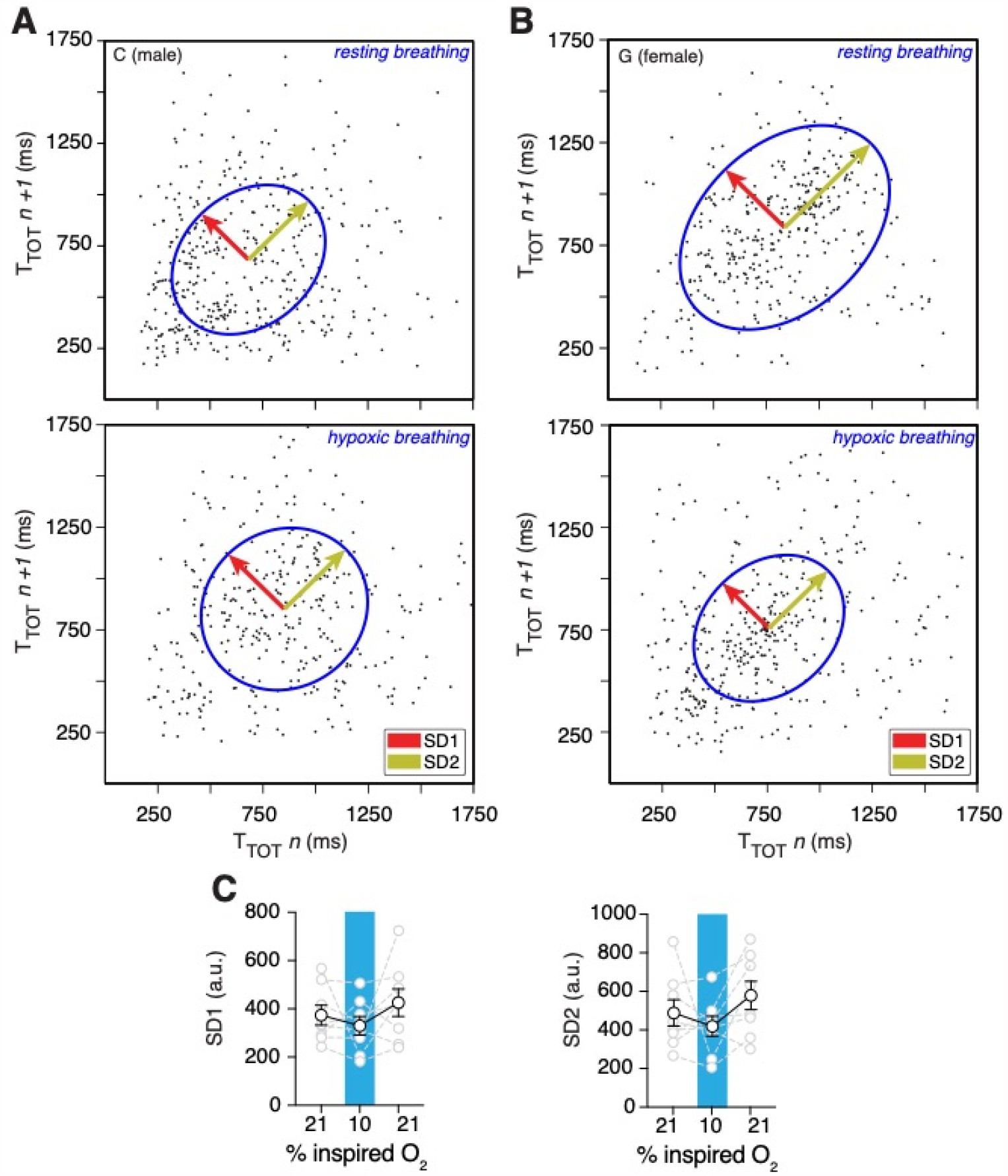
Changes in respiratory rate variability following hypoxia challenge. Representative Poincaré plots of total cycle duration (T_TOT_) for the n^th^ cycle versus T_TOT_ for the n^th^+1 cycle during baseline and hypoxic conditions (10% O_2_) in male (**A**) and female (**B**) marmosets. (**C**) Summary data illustrating changes in SD1 and SD2 before, during, and after hypoxic challenge. Respiratory rate variability did not change for either measure during (SD1: *p* = 0.4; SD2: *p* = 0.6; Wilcoxon matched-pairs signed rank test) or after (SD1: *p* = 0.6; SD2: *p* = 0.3; Wilcoxon matched-pairs signed rank test) the challenge compared to baseline. Data are shown as individual (gray lines) and mean (black line) values ± SEM. a. u. – arbitrary unit.

We then calculated the metabolic rate (M_R_) in marmosets during hypoxic challenge. Our data suggest that M_R_ had a profound decrease (∼ 50%) during hypoxia when compared to the baseline (Table 1).

**Table 1.** Hypoxia decreased metabolic rate in common marmoset.

|  | *pre-hypoxia* | *hypoxia* | *post-hypoxia* |
| --- | --- | --- | --- |
| Metabolic rate (%) | 100 | 51 ± 4 | 98 ± 1 |

### Post-Hypoxic Ventilatory Response

We also measured changes in respiratory features in the five minutes immediately following the hypoxic challenge (post-hypoxic challenge). Though we saw no changes in *f*_*R*_ (73 ± 4 vs 82 ± 7 breaths min^-1^ in baseline, *p* = 0.11, Wilcoxon matched-pairs signed rank test) and *V*_T_ (0.47 ± .10 vs 0.54 ± 0.12 a.u. in baseline *p* = 0.15, Wilcoxon matched-pairs signed rank test), *V*_E_ decreased (34 ± 7 vs 46 ± 12 a.u. in baseline, *p* = 0.078, Wilcoxon matched-pairs signed rank test) relative to baseline (Figure 6).

We also calculate changes in T_I_, T_E_, and R_D_, immediately following the hypoxia challenge. While we observed no change in T_E_ (5.4 ± 0.5 vs 5.0 ± 0.6 a.u. in baseline, *p* = 0.46), there was an increase in T_I_ (3.4 ± 0.2 vs 3.0 ± 0.2 a.u. in baseline, *p* = 0.055) and a decrease in R_D_ (1.3 ± 0.2 vs 1.8 ± 0.4 a.u. in baseline, *p* = 0.078) after hypoxia challenge (Figure 7).

### Sigh frequency, sniffing, and apnea index

Since incidences of sighs, apneas, and sniffing could contribute to the irregularity of respiration, we measured frequencies of these essential features of breathing behavior. It has been shown that sigh can be generated within the inspiratory rhythm-generating circuits of the preBötC (Sheikhbahaei et al. 2018; Li et al. 2016; Lieske et al. 2000; Borrus et al. 2020; Toporikova et al. 2015; Vlemincx et al. 2013), and may be modulated by excitatory signals from central chemocenters (Sheikhbahaei et al. 2018; Sheikhbahaei et al. 2017; Souza et al. 2018; Souza et al. 2019; Li et al. 2016). In female adult marmosets, sigh frequencies were not different when compared to those in male animals during the baseline in room air (11 ± 1 vs. 12 ± 2 hr^-1^ in male) (Figure 9 – figure supplement 1A). Both hypoxic and hypercapnic challenges increased frequency of sighs in rodents (Li et al. 2016; Sheikhbahaei et al. 2018). Consistent with those results, hypoxia increased sigh events by 5.5 folds in marmoset (71 ± 10 vs. 11 ± 1 hr^-1^ in room air, *p* = 0.008, Wilcoxon matched-pairs signed rank test). Similarly, hypercapnia also increased sigh frequency (68 ± 3 vs. 12 ± 1 hr^-1^ in room air; *p* = 0.008, Wilcoxon matched-pairs signed rank test) (Figure 9A)

**Figure 9.**
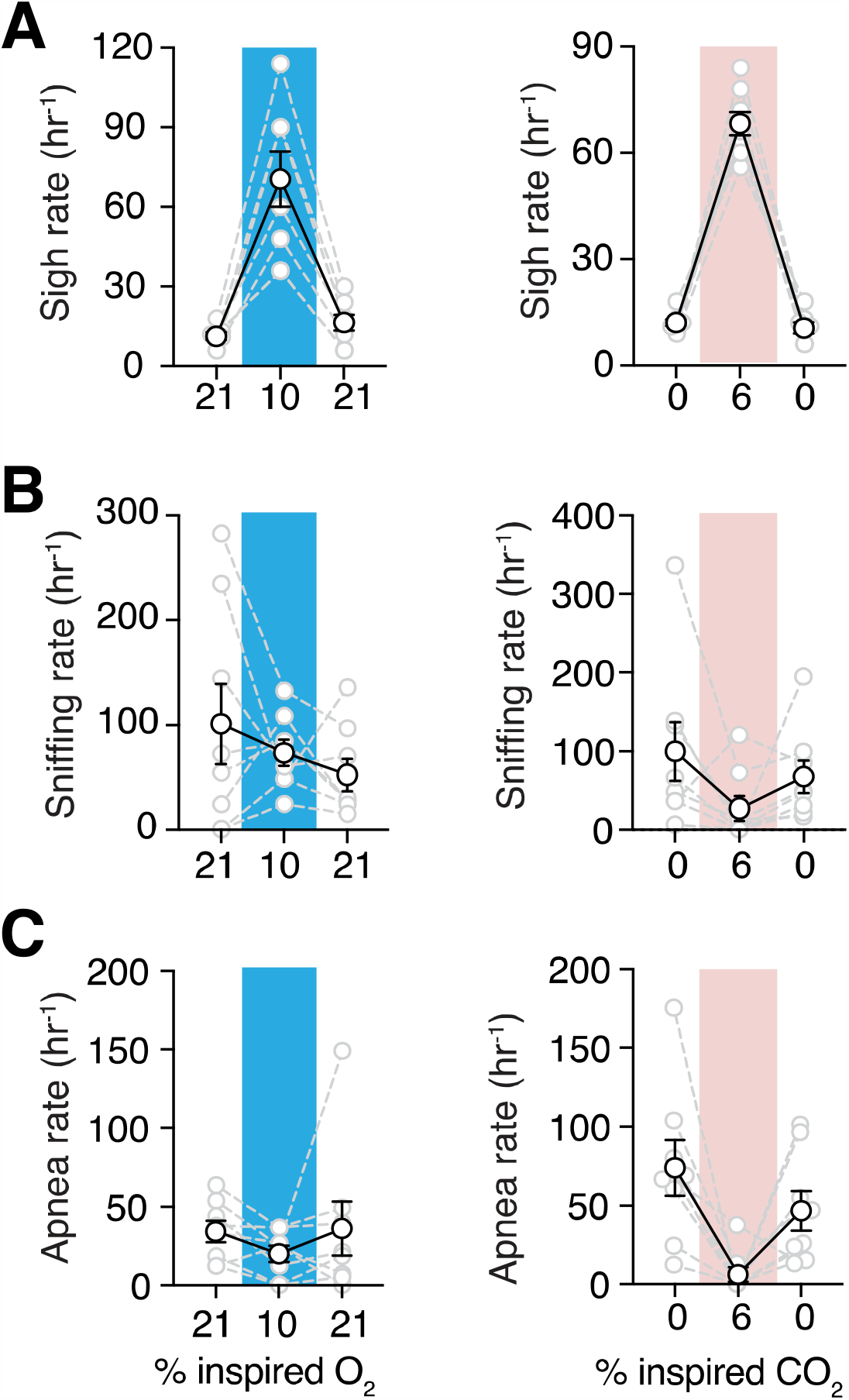
Sigh frequencies, sniffing rate, and apnea index during hypoxia and hypercapnia challenges. (**A**) Summary data demonstrating increase in sigh frequency after 5 minutes of hypoxic (10% O2, *left*) or hypercapnic (6% CO2, *right*) challenge (*p* = 0.008 and *p* = 0.008, respectively; Wilcoxon matched-pairs signed rank test). (**B**) Summary data demonstrating no change in sniffing rate during (*p* = 0.74) and after (*p* = 0.74) hypoxia challenge (*left*). Sniffing rate increased during and returned to baseline after hypercapnia challenge (*p* = 0.008 and *p* = 0.08 respectively; Wilcoxon matched-pairs signed rank test) (*right*). (**C**) Grouped data demonstrating decrease in rate of spontaneous apneas during hypoxia (*p* = 0.04, Wilcoxon matched-pairs signed rank test) (*left*) and hypercapnia (*p* = 0.008, Wilcoxon matched-pairs signed rank test) (*right*). Data are shown as individual (gray lines) and mean (black line) values ± SEM.

We also analyzed high frequency breathing (sniffing) in marmosets. During hypoxic challenge, the sniffing rate did not change with respect to baseline (74 ± 13 vs. 101 ± 38 hr^-1^ in baseline, *p* = 0.84, Wilcoxon matched-pairs signed rank test) (Figure 9B). However, during hypercapnic challenge, rate of sniffing was less than that in room air (27 ± 16 vs. 100 ± 38 hr^-1^ in baseline, *p* = 0.078, Wilcoxon matched-pairs signed rank test) (Figure 9B).

Spontaneous and post-sigh apneas are reported in rodents, rabbits, humans, and other animals (Yamauchi et al. 2008; Franco et al. 2003; van der Heijden and Zoghbi 2018; Bongianni et al. 2010; Li et al. 2006; Ramirez et al. 2013; Sheikhbahaei et al. 2017). We did not find differences in apnea index between female and male marmosets (Figure 9 – figure supplement 1C). Apneas decreased during hypoxic challenge (37 ± 12 vs. 79 ± 20 in room air, *p* = 0.039, Wilcoxon matched-pairs signed rank test) (Figure 9C). During hypercapnic challenge, rate of spontaneous apneas also decreased drastically relative to that in room air (9 ± 6 vs 129 ± 31 hr^-1^ in room air, *p* = 0.008, Wilcoxon matched-pairs signed rank test) (Figure 9C).

### Spontaneous activity

Lastly, we measured the activity of marmosets in the plethysmograph during hypoxic and hypercapnic challenges. In measurements of large changes in position from one quadrant of the chamber to another, we saw no changes in hypoxia challenge (4.1 ± 1.8 vs 5.4 ± 1.6 quadrant changes per minute at baseline, *n* = 3, *p* = 0.99, Wilcoxon matched-pairs signed rank test) or hypercapnia challenge (4.0 ± 0.5 vs 4.7 ± 1.4 quadrant changes per minute at baseline, *n* = 3, *p* = 0.75, Wilcoxon matched-pairs signed rank test). Similarly, we observed no differences in sum of frame-to-frame Euclidean distances in hypoxia challenge (124 ± 28 vs 128 ± 27 pixels min^-1^ at baseline, *n* = 3, *p* = 0.99, Wilcoxon matched-pairs signed rank test) or hypercapnia challenge (126 ± 20 vs 120 ± 11 Px /min at baseline, *p* = 0.99, Wilcoxon matched-pairs signed rank test) (Figure 10)

**Figure 10.**
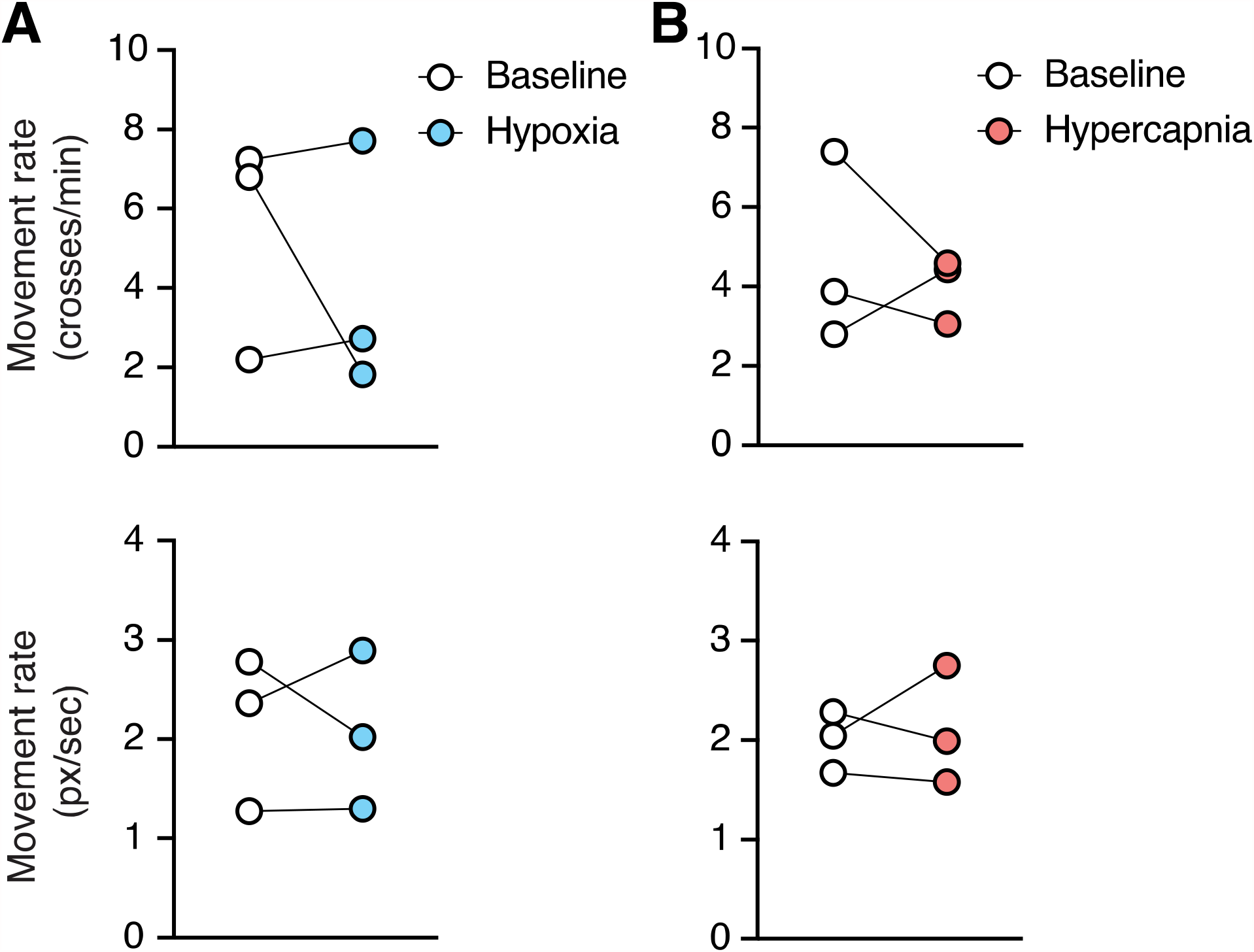
Changes in spontaneous activity during hypoxic and hypercapnic challenges. (**A**) Hypoxia did not induce any changes in animal movement rate as measured by quadrant changes in the chamber (*top*) (*p* = 0.99, *n* = 3; Wilcoxon matched-pairs signed rank test), or as measured by total change in animal position per second (*bottom*) (*p* = 0.99, *n* = 3; Wilcoxon matched-pairs signed rank test). (**B**) We detected no changes in animal’s movement rate as measured by quadrant changes in the chamber (*top*) (*p* = 0.75, *n* = 3; Wilcoxon matched-pairs signed rank test), or by total change in position per second (*bottom*) (*p* = 0.99, *n* = 3; Wilcoxon matched-pairs signed rank test) during hypercapnia.

## Discussions

We used non-invasive, Whole-body Plethysmography to measure breathing behaviors (Hamelmann et al. 1997) in unrestrained, freely moving, awake marmosets. Plethysmography has a simple and robust design that has been used widely in humans [neonates (Sivieri et al. 2017) and adults (Dubois et al. 1956)], non-human primates [such as macaques (Besch et al. 1996) and cynomolgus monkeys (Iizuka et al. 2010)], rodents (Sheikhbahaei et al. 2018; Hosford et al. 2020), dogs (Liu et al. 2016), sheep (Hutchison et al. 1983), cats (Hoffman et al. 1999), turtles (Valente et al. 2012), and other animals.

The common marmoset (*Callithrix jacchus*) is a small New World primate (Okano et al. 2015). Recently, marmosets have been proposed as a powerful animal model in neuroscience research (Miller et al. 2016; Burkart and Finkenwirth 2015; Leopold et al. 2017; Mitchell and Leopold 2015), especially to study vocal communication (Eliades and Miller 2017). Compared to rodents, marmoset’s central nervous system more closely resemble humans’ in terms of physiological function and anatomy of the brain (Bendor and Wang 2005). In addition, considering the similarity of the brain structure and circuit connectivity between primates, marmosets provide an attractive opportunity to study cortical (i.e., voluntary) control of motor activity (Walker et al. 2017) as well as coordination of breathing with complex functions, such as vocalization. However, the basic characteristics of breathing behaviors in common marmoset were not yet defined. Accordingly, in this study we characterize respiratory behaviors in awake, freely behaving marmosets in their natural posture. Using the whole-body respiratory measurement in conscious animals requires sophisticated algorithms to distinguish the respiratory signals from noises (i.e., movements). To avoid this problem, respiratory activities are often recorded when the animal is asleep or anesthetized. There are absolute advantages to studying the homeostatic control of breathing physiology in awake animals, despite the increased variability. Therefore, to overcome this challenging task, we validated and optimized a new Python package, Neurokit2 (see Methods section), for unsupervised analysis of respiratory signals obtained from experimental animals. We then characterized the ventilatory responses to hypoxia (decrease inspired O_2_ to 10%) and hypercapnia (increased inspired CO_2_ to 6%).

### The hypercapnic ventilatory response

Currently, the chemosensitivity mechanisms that adjust breathing with respect to the level of *P*CO_2_ / pH in the brain are centered around neurons and astrocytes in the retrotrapezoid nucleus (RTN) and medullary raphé (Kumar et al. 2015; Teran et al. 2014; Guyenet et al. 2019). However, other data support a hypothesis that distributed chemosensitive regions in the medulla act as central respiratory chemosensors and are responsible for mounting of about 70% of the hypercapnic respiratory response (the mechanism that adjusts breathing in accordance with increase in *P*CO_2_) (Nattie 1999; Nattie 2000; Nattie 2001; Spyer and Thomas 2000; Nattie and Li 2009). Specialized peripheral chemoreceptors located in the carotid bodies (and aortic bodies in some species) are responsible to the remaining 30% of hypercapnia-induced augmentation of breathing. In our experiments, to prevent hypoxia, we applied hyperoxic hypercapnia (60% O_2_ / 6% CO_2_ balanced with N_2_) as it is routinely used in rodent experiments (Teppema et al. 1997; Hermand et al. 2015; Sheikhbahaei et al. 2018; Hosford et al. 2020). It has been shown that hyperoxia minimizes the input from peripheral chemosensors of carotid bodies (Chavez-Valdez et al. 2012; Gonzalez et al. 1994). Since marmosets lack aortic bodies (Clarke and de B Daly 2002), the hypercapnic ventilatory response reported here is driven by the central CO_2_ respiratory chemocenters. In awake, freely behaving marmosets, hypercapnia increased both breathing rate (*f*_R_) and tidal volume (*V*_T_) (Figure 3). However, the augmentation of ventilation (*V*_E_) was mainly due to increase in *V*_T_ (by ∼ 160%) rather than *f*_R_. These data are comparable to data from human (Duffin et al. 2000) as well as recent data obtained from rodents (Bhandare et al. 2020; Sheikhbahaei et al. 2018). Hypercapnia also increases frequency of sighs (i.e., augmented breath) in rodent (Forsberg et al. 2016; Ramirez 2014). Consistent with these data, we also found that hypercapnia increased sigh frequency in marmosets (Figure 9).

### The hypoxic ventilatory response

The hypoxic ventilatory response (HVR) in common marmoset was interesting as there was little or no increase in *f*_R_ and *V*_T_ during hypoxic exposure (Figure 6). We believe the level of O_2_ during hypoxia was sufficient to elicit HVR as similar level of O_2_ (10% O_2_ for 180 seconds) decreased the peripheral oxygen saturation (SpO_2_) to 89% in humans (Gerlach et al. 2021). In addition, increases in sigh rate, existence of post-hypoxic depression (see below), and decrease in metabolic rate (M_R_) strongly suggests that the respiratory circuits were activated by the hypoxic challenge to prevent hypoxic ventilatory decline (HVD). It is possible that large gas-exchange capacity of marmosets (due to the increased oxygen diffusion capacity of their lungs) (Barbier and Bachofen 2000) maintains the adequate blood oxygenation, and therefore, blunt the HVR.

Although hypoxic conditions in marmoset’s habitat (sea-level forests of the Amazon) are rare, hypoxia might occur during sleep (i.e., sleep apnea) or as a result of a disease state. It is commonly believed that the HVR is biphasic in adult mammals. During acute hypoxia, ventilation is depicted by an initial increase followed by a subsequent decline to a value at or above the baseline (i.e., HVD). This biphasic hypoxic response has been reported in humans, rats, and other mammals (Eden and Hanson 1987; Martin et al. 1990; Fung et al. 1996; Dahan et al. 1996; Vizek and Bonora 1998), however, earlier reports suggest that there are considerable interindividual variations in HVR in humans (Hirshman et al. 1975; Weil and Zwillich 1976). In awake adult human, even one-hour hypoxic condition did not affect mean pulmonary minute ventilation (i.e., *V*_E_) (Robinson and Haymes 1990; Seo et al. 2017). Similarly, recent data in awake adult humans showed no increase of *f*_R_ during acute hypoxia (Gerlach et al. 2021), suggesting that the any changes in ventilation may be due to increase in *V*_T_, not *f*_R_ (Tarbichi et al. 2003). Our data agree with these reports as we see variable responses to hypoxia in marmosets, absence of hypoxic-induced increase in *V*_E_, as well as a slight increase in *V*_T_ and *V*_E_ during the first minute of HVR (Figure 6).

Hypoxia increases sigh frequency in mammals, even in animals whose carotid bodies are not functional (Bartlett 1971; Schwenke and Cragg 2000; Cherniack et al. 1981; Sheikhbahaei et al. 2018). Consistent with other data, hypoxia also increased sigh frequency in marmosets. In addition, the fact that sigh frequency, but not the breathing frequency, increased during hypoxic challenge supports the hypothesis that distinct cells (possibly within the preBötC region) may be responsible for generation of rhythmic sigh and normal breathing (Toporikova et al. 2015; Li et al. 2016; Sheikhbahaei et al. 2018). Recent data from behaving rats suggest that purinergic signaling from preBötC astrocytes may play a significant role in sigh generation (Sheikhbahaei et al. 2018).

On the other hand, the mechanism of hypoxic ventilatory decline (HVD) is not fully understood. It is proposed that desensitization of peripheral chemoreceptors might have a role (Bascom et al. 1990), though significant evidence suggest that, at least in rodents, astrocytes (the numerous star-shaped glia cells) in the preBötC are capable of acting as central respiratory oxygen chemosensors (Sheikhbahaei et al. 2018; Angelova et al. 2015; Rajani et al. 2018) and contribute to the HVD possibly via release of adenosine triphosphate (ATP) (Sheikhbahaei et al. 2018). Existence of ATP receptors in the brainstem respiratory nuclei of marmosets (Yao et al. 2000) further strengthens this hypothesis in primates. In addition to preBötC, retrotrapezoid nucleus (RTN), rostral ventrolateral medulla (rVLM) and the nucleus of the solitary tract (NTS) in the brainstem are proposed to have oxygen sensing capabilities (Accorsi-Mendonça et al. 2015; Mazza et al. 2000; Uchiyama et al. 2020). However, more research is required to understand if this ‘distributed central oxygen chemosensors’ hypothesis (Sheikhbahaei 2020) can be generalized to primates.

Decrease of post-hypoxic ventilation in human is also reported (Tarbichi et al. 2003). Similarly, we observe such a respiratory response in marmoset. This post-hypoxic depression is also illustrated in conscious (Angelova et al. 2015; Sheikhbahaei et al. 2018) and anesthetized (Rajani et al. 2018) rats. However, we did not detect any sex differences during post-hypoxic recovery from hypoxia as reported in rat’s *in vitro* models (Garcia et al. 2013). Our data is consistent with the data reported in human that there is no differences in ventilation between the sexes during post-hypoxic response (Tarbichi et al. 2003).

Other than an increase in ventilatory response during hypoxia, mammals can reduce oxygen demand by optimizing and decreasing their rates of metabolism (Dzal et al. 2015). During hypoxia, adult marmosets decreased their metabolic rates by ∼ 50%, which is similar to the data reported in other primates [pygmy marmosets (Tattersall et al. 2002), and human (Robinson and Haymes 1990; Knuttgen and Saltin 1973; Consolazio et al. 1966; Adams and Welch 1980)], but 2-3 times more than the calculated rates from cats (Gautier et al. 1989) and rats (Mortola et al. 1994). This decrease in metabolism together with increase in sigh frequency might be sufficient for homeostatic control of blood oxygen during acute hypoxia in primates. We also looked at marmoset activity during hypoxic challenges. However, our data suggest that animals’ activity were not affected by hypoxia. This suggests that decreases in metabolic rate during hypoxic challenge are not due to decrease in spontaneous activity, but may be accounted for by changes in other processes with metabolic demand such as thermoregulation or cardiovascular activities.

The brain is highly susceptible to low oxygen levels. Supply of oxygen may be decreased in clinical conditions (such as sleep apnea or stroke) or environmental settings (such as exposure to carbon monoxide). Therefore, an understanding of how the brain maintains homeostatic levels of oxygen and responds to hypoxic events is of longstanding interest. Studies in common marmosets will shed some light on this problem and might fill the gap between rodent and human research to better understand the homeostatic control of breathing and its disorders.

## Acknowledgements

This work was supported by the Intramural Research Program (IRP) of the NIH, NINDS and NIMH. We are grateful for invaluable supports and discussions from Drs. David Leopold, Yogita Chudasama, and Jeffrey Smith. We also thank Dr. Gregory Funk for valuable consultations.

## Notes

<u>Conflict of interest:</u> The authors declare no competing financial interests.

https://github.com/NGSC-NINDS/Marm_Breathing_Bishop_et_al_2021

### Competing Interest Statement

The authors have declared no competing interest.

